# Cross-sex cecal microbiota transfer alters depressive-like behaviours in mice

**DOI:** 10.1101/2022.10.12.511923

**Authors:** Meagan Hinks, Yellow Martin, Francine Burke, Francis R. Bambico, Ashlyn Swift-Gallant

## Abstract

Major depressive disorder (MDD) is a leading cause of non-fatal global disease burden, with females being two-fold more likely than males to be diagnosed with the disorder. Despite this distinct sex-linked disparity of diagnosis, it is unclear what underlies the sex bias of MDD prevalence. Recent findings suggest a role for the gut in mediating affective disorders through the gut-brain-axis (GBA). However, few studies have included sex as a biological variable. For this study, cross-sex transfer of cecal microbiota was performed between male and female C57Bl/6 mice to elucidate the effects of sex and the gut microbiome on a standard battery of tests measuring depressive-like behaviours. Specifically, regardless of sex, recipients of male cecal content had a greater sucrose preference than controls and recipients of female cecum. Conversely, in the splash test, recipients of female cecum displayed a decrease in grooming behaviour compared to both controls and recipients of male cecum, suggestive of an increase in depressive-like behaviour. These results support a role for female-specific gut microbes in contributing to female vulnerability to depression, while male-specific gut microbes may protect in part against an anhedonia-like phenotype

---

Major depressive disorder (MDD) is one of the leading causes of non-fatal illness worldwide [1]. Among those affected, females are two times more likely to be diagnosed with MDD than males [1,2]. Emerging evidence suggests that the gut may contribute to susceptibility of MDD via communication with the brain, also known as the gut-brain axis (GBA) [3,4,5]. As gut microbial composition has been found to vary by sex, sex-linked differences in composition may be a source for the apparent sex-disparity in diagnosis of MDD [6].

Various studies have investigated the impact of the gut microbiome on depressive-like symptoms using rodent models with various microbial altered strains (i.e., germ-free (GF) and specific-pathogen free (SPF) mice) as well as fecal/cecal transplantation from depressed patients or mouse models of depression to healthy rodents [5,7,8,9]. For example, mice void of any microorganisms (i.e., GF mice) exhibit decreases in behavioural stress-response than SPF mice or controls [5,8,9]. Furthermore, fecal/cecal transplantation from either human patients or mouse models of depression to otherwise healthy mice is sufficient to induce a depressive-like phenotype [4,5,7,9). However, such studies have strictly used male mouse models and/or do not identify the sex of recipients/donors [4,5,7,9]. Given sex differences in both gut microbiome composition and in susceptibility of depression, it is critical to include sex as a biological variable to understand whether the role of the gut for depression is also sex dependent.

In the present study, we assessed whether cross-sex cecal transfer would be sufficient to alter behaviour on a standard battery of tests for a depression-like phenotype. Since females present a more pronounced susceptibility to MDD, we predicted that female cecal transplantation would result in an increased depressive-like phenotype in the receivers, regardless of the sex of the recipient. Conversely, we predicted transfer of male cecal content would decrease depressive-like behaviours in recipients.

To address this question, we obtained mice from Charles River Laboratories (Quebec, Canada). Mice were singly housed, provided *ad libitum* access to water and food (Harlan Laboratory Chow) and kept on a 12:12 light/dark cycle. Sixty C57BL/6 mice, half male, half female, were assigned to one of three treatment groups: vehicle (saline control), female cecum, or male cecum (n = 10 males and 10 females per treatment group). Cecal treatments began between postnatal day (PND) 60-95, and cecum was administered via oral gavage using a 20-gauge ball-tipped gavage needle. Mice were treated once every other day for 12 days, for a total of six treatments. Mice received 0.2mL of either cecum (diluted 1:10 with saline) or saline vehicle on treatment days. Ethical approval for this study was obtained from the Institutional Animal Care Committee at Memorial University of Newfoundland and all procedures complied with the Canadian Council of Animal Care (CCAC) guidelines.

Cecum donor mice were also obtained from Charles River under the same housing conditions (n = 6-8/sex). Donor mice were overdosed with Avertin (40mg/kg) via intraperitoneal injection in the lower right abdominal quadrant to prevent injection directly into the cecum. An incision was then made medially down the abdomen, the cecum sac removed from the intestinal tract and the cecal contents were collected in 1.7ml microcentrifuge tubes, combined by sex. Cecum was diluted 1:1 with 0.1M phosphate buffered saline (PBS) prior to storage. Cecum was stored at -80 degrees Celsius, and aliquots were thawed on the day of treatment, and diluted 1:5 with saline for a total dilution of 1:10.

## Behavioural Testing

Following cecal/saline treatment, mice were assessed for depressive-like behaviours on a behavioural test battery. This battery comprised of 4 tests for depressive-like symptomology: sucrose preference, splash, tail suspension, and forced swim tests. Following behaviour testing, mice were weighed prior to euthanasia to assess whether weights varied by sex and/or between treatment groups. Weights and behavioural indices of depression were analyzed for sex by treatment effects using between-subjects analysis of variance (ANOVA) followed by planned comparisons for significant effects (alpha set at *p* < .05).

> ***Sucrose Preference Test:*** The sucrose preference test (SPT) is a measure used to assess anhedonia, the decreased ability to experience pleasure, one of the core symptoms of depression [10]. For this test, mice were first habituated to drinking water from two small plastic cups in their home cages for 18-24h. After the habituation phase, one cup was filled with regular drinking water and the other cup filled with a 4% sucrose solution.
>
> Following 18 hours, the weight of each cup was recorded and deducted from the starting weight. A preference score was then recorded by subtracting the weight of the water drank from the weight of the sucrose solution. A high sucrose preference criterion was set at 70% based on typical cut-off values which vary between 65-80% [11]. Data from this test were omitted for two female mice whose cups had been spilled during the testing phase.
>
> ***Splash Test*:** In the splash test (ST), a single mouse was placed in an empty mouse cage and provided with five minutes to habituate to the new environment. Then, a viscous 20% sucrose solution was applied to the dorsal coat of the mouse. The test was then video recorded for five minutes and later scored for grooming and general locomotor behaviours. Grooming behaviour was used as an indication of self-care and motivational behaviour, while decreased time spent grooming indicates depressive-like behaviour [12].
>
> ***Tail Suspension Test***: In the tail suspension test mice were suspended by their tail, close to its base (i.e., tail is taped to the top of a rodent cage) so that the body dangles, facing downward for five minutes. Each trial was video recorded, with immobility (measure of depressive-like behaviour) and escape behaviours (indication of motivational behaviour) measured and analyzed for both duration and frequency with Ethovision XT14 Software. Escape behaviours consist of movements of the hinds/forelimbs as well as increased number of attempts/min to reach their tail [13].
>
> ***Forced Swim Test***: Mice were placed in a transparent glass cylinder (38.0 cm height x 27.0 cm diameter) filled with water (at a temperature of 25°C +/-0.5°C). The mice could not reach the top or bottom of the cylinder, and thus are suspended in the water for the duration of the 5-minute test. The video recordings of the trials were then analysed for duration and frequency of depressive-like behaviour (i.e., immobility) using Ethovision XT14 software [14].

Our results suggest there are sex-specific effects of the gut microbiome on depressive-like behaviour. Specifically, regardless of sex, recipients of male cecal content had a greater sucrose preference than controls, and a higher proportion of mice receiving male cecum were classified as having high preference scores (>70%) than both controls and recipients of female cecum (Figure 1). In the splash test, female cecum decreased grooming behaviour in recipients compared to both controls and recipients of male cecum, suggestive of an increase in depressive-like behaviour. While the omnibus effect was only trending (p = .076), results on the forced swim test suggest that male cecum increased immobility, indicative of depressive-like phenotype, compared to control mice (posthoc *p* = .032, *d* = 0.7, Figure 1). No significant differences in general locomotor behaviour was found on the splash test, indicating that increased immobility on the forced swim test was due to passive coping, not general differences in locomotor activity. No differences found on tail suspension and no effects of treatment were found on body weight at the end of the experiment. Though we did find the expected effect of sex, such that male mice had a greater body weight (*M* = 26.6, *SEM* = 0.351) than females (*M* = 21.4, *SEM* = 0.358), *F* (2,54) = 102.657, *p* ***<*** .001.

**Figure 1.**
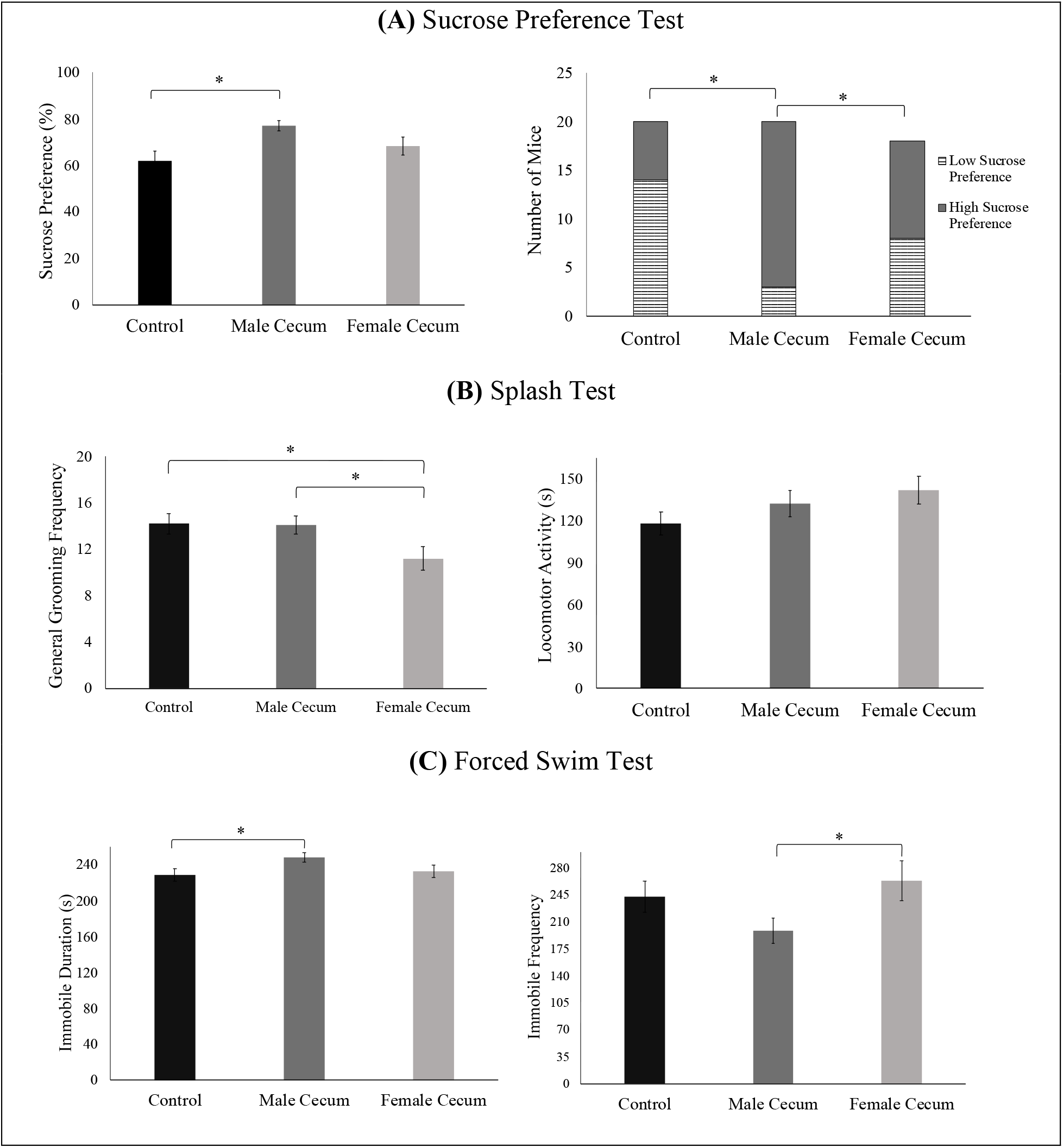
Data for behavioural indices of depressive-like behaviour on the sucrose preference, splash, and forced swim tests. **Note**. Data shown as mean ± standard error of the mean (SEM) for (**A**) sucrose preference, (**B**) splash, and (**C**) forced swim tests. **A**. Mice treated with male cecum had significantly higher sucrose preference scores compared to controls (*F* (2,52) = 5.154, *p* ***=*** .009, post hoc *p* = .002, Cohen’s *d* = 1.01), as well as a significantly higher proportion of mice classified as having high preference scores compared to the control and female cecum groups. Chi-square analysis of the proportion of mice who had a high sucrose preference score (≥70%) compared to a low score (<70%) showed a significant main effect of treatment, *X*^*2*^ (2, *N =*58) *=* 12.534, *p* **=** .002, Cramer’s *V* = 0.462. A follow-up series of chi-square tests revealed that the proportion of mice in the male cecum condition with a high sucrose preference score was significantly greater (*N* = 17/20) than the proportion of those in the female cecum condition (*N* = 10/18), *X*^*2*^ (1, *N =*38) *=* 3.993, *p* = .046, *OR* = 4.533, 95% CI [0.972, 21.141] and control group (*N* = 6/20), *X*^*2*^ (1, *N =*40) *=* 12.379, *p* < .001, *OR* = 13.222, 95% CI [2.790, 62.670]). **B**. Mice treated with female cecum had significantly *lower* grooming frequenc*y* compared to *both* controls *and recipients of male cecum* (*F* (2,51) = 3.949, *p* = .025, post hoc *p* = .032, *d* = 0.73 and *p* = .011, *d* = 0.86, respectively). No differences in general locomotor activity were found, *p >* .*05*. **C**. Mice treated with male cecum had significantly lower immobile frequencies compared to mice treated with female cecum *(F* (2,54) = 2.464, *p =* .095, post hoc *p* = .034, *d* = 0.69) and significantly higher immobile durations compared to controls *(F* (2,54) = 2.701, *p =* .076, post hoc *p* = .032, *d* = 0.7). *\* indicates p <* .*05*.

Overall, our findings suggest that female cecal content increases depressive-like behaviour on measures of self-care/motivational behaviours, while male cecum increases hedonic behaviours as measured by the sucrose preference test as well leading to decreased immobility on the force swim, a measure of passive coping behaviour. These results suggest that the effects of the gut microbiome on depressive-like behaviours are: 1) not entirely due to depression itself; cecal transfer from healthy animals can change depression-like behaviours, and thus it may be that gut microbes regulate mood more broadly, rather than depression specifically. However, these regulators of mood may also contribute to the sex bias in susceptibility to affective disorders. Thus, sex differences within the gut microbiome in healthy/unmanipulated mice may contribute to vulnerability to depressive-like behaviours, as previously suggested [15,16,17]. These results also suggest, 2) there are distinctions in the role of the gut microbiome/sex for self-care vs escape behaviours, such that female-typical gut composition may decrease self-care/motivational behaviours while male-typical gut composition may decrease escape behaviours. These findings are consistent with sex differences in these behaviours. For examples females tend to exhibit escape behaviours in response to stressors, while males tend to show immobility [18,19], and thus our findings suggest that the gut microbiome mediates, at least in part, sex differences in these behaviours.

## Conclusion

We found support for our predictions that the female gut microbiota increases while male microbiota decreases depressive-like behaviours. Unexpectedly, we found that male gut microbiota increased depressive-like behaviour on measures of coping behaviours (i.e., the forced swim test). Further work is needed to delineate which microbes mediate these effects, and whether they act directly via communication with the brain (i.e., vagus nerve) or indirectly via mediation of neuroendocrine and/or immune factors [20]. Nevertheless, these results suggest that there are sex-specific gut microbes which mediate behaviours on the standard battery of tests for depression in rodent models. Thus, sex differences on measures of depression may be due in part to sex differences in gut microbiota composition, and these sex differences in gut microbes may underlie the sex bias in vulnerability to depression. This study provides a basis for the consideration of sex and its relationship to depression and the GBA using mouse models of depressive-like behaviour.

